# Dermatan Sulfate Is a Potential Master Regulator of IgH via Interactions with Pre-BCR, GTF2I, and BiP ER Complex in Pre-B Lymphoblasts

**DOI:** 10.1101/2021.01.18.427153

**Authors:** Jongmin Lee, Jung-hyun Rho, Michael H. Roehrl, Julia Y. Wang

## Abstract

Dermatan sulfate (DS) and autoantigen (autoAg) complexes are capable of stimulating autoreactive CD5+ B-1 cells. We examined the activity of DS on CD5+ pre-B lymphoblast NFS-25 cells. CD19, CD5, CD72, PI3K, and Fas possess varying degrees of DS affinity. The three pre-BCR components, Ig heavy chain mu (IgH), VpreB, and lambda 5, display differential DS affinities, with IgH having the strongest affinity. DS attaches to NFS-25 cells, gradually accumulates in the ER, and eventually localizes to the nucleus. DS and IgH co-localize on the cell surface and in the ER. DS associates strongly with 17 ER proteins (e.g., BiP/Grp78, Grp94, Hsp90ab1, Ganab, Vcp, Canx, Kpnb1, Prkcsh, Pdia3), which points to an IgH-associated multiprotein complex in the ER. In addition, DS interacts with nuclear proteins (Ncl, Xrcc6, Prmt5, Eftud2, Supt16h) and Lck. We also discovered that DS binds GTF2I, a required gene transcription factor at the *IgH* locus. These findings support DS as a potential master regulator of IgH in pre-B cells at protein and gene levels. We propose a (DS•autoAg)-autoBCR dual signal model in which an autoBCR is engaged by both autoAg and DS, and, once internalized, DS recruits a cascade of molecules that may help avert apoptosis and steer autoreactive B cell fate. Through its affinity with autoAgs and its control of IgH, DS emerges as a potential key player in the development of autoreactive B cells and autoimmunity.

## Introduction

Autoimmunity, an immune response against the body self, is an intriguing phenomenon. Autoimmunity is caused by autoreactive lymphocytes and/or autoantibodies that target autoantigens (autoAgs) present in single or multiple tissue types throughout the body, leading to over 80 recognized types of autoimmune diseases, ranging from systemic diseases such as lupus and rheumatoid arthritis to organ-specific diseases such as type 1 diabetes. Autoimmune phenomena are also strongly linked to various malignancies such as lymphoma and leukemia. Despite extensive research, understanding how autoreactive lymphocytes survive rounds of immune editing and elimination has remained largely elusive. Although it is now accepted that autoreactive cells are positively selected by autoAgs, the underlying mechanism is unclear. Furthermore, several hundred autoAgs have been identified across seemingly unrelated tissue locations or biologic functions. Yet these different autoantigens can all elicit a similar autoimmune response such as clonal expansion of autoreactive cells or production of specific autoantibodies. Better understanding of these processes will greatly advance the field of autoimmunity.

We have proposed that the glycosaminoglycan dermatan sulfate (DS) is a key missing player in autoimmunity in that (i) self-molecules with affinity to DS have a high propensity to be autoAgs and (ii) DS•autoAg complexes work in concert to stimulate autoreactive B cells (*1, 2*). In our initial study, we had injected mice with various glycosaminoglycans, including DS, heparan sulfate, chondroitin sulfates A and C, hyaluronic acid, and heparin, and found that DS is the most potent in inducing arthritis in mice (*3*). When tested in vitro with mouse splenocytes, DS is found to be the most potent in stimulating CD5+ B (B-1) cell proliferation and, moreover, DS exerts unique affinity to autoantigens from apoptotic cells, which prompted our hypothesis that DS and autoAgs form complexes and co-operate to stimulate B-1 cells (*1*). We then tested clinical autoantibody standard sera and sera from patients with autoimmune diseases such as lupus and Sjögren syndrome and demonstrated that DS affinity enabled the identification of many more autoAgs than current clinical tests are able to (*2*). Based on our DS•autoAg affinity hypothesis, we have thus far identified over 200 potential protein autoAgs from cells lines and murine liver and kidney tissues (*2, 4, 5*).

When autoAg•DS complexes encounter B cells expressing autoreactive B cell receptors (autoBCRs), we hypothesize that, while autoAgs engage autoBCRs, DS must engage other signals to ensure successful B-1 cell activation leading to further proliferation and differentiation. Neither autoAgs nor DS alone can recruit a sufficient signaling network, as autoAg-autoBCR engagement without co-stimulating factors generally leads to deletion or anergy of the autoreactive cells. In our search for DS receptor(s) in autoreactive B-1 cells, we investigated CD5+ pre-B lymphoblast NFS-25 cells, a cell line derived from a spontaneous murine lymphoma (*6*). The pre-B stage is a crucial checkpoint for positive vs. negative regulation of autoimmunity and for development of mature autoreactive B cells. Precursor BCRs (pre-BCRs) are considered poly- and self-reactive, resembling polyreactive autoBCRs (*7*). Pre-BCR-mediated signaling has been implicated in positive selection of pre-B cells, proliferation, survival, IgH allelic exclusion, and IgH repertoire selection (*8*). The pre-BCR is composed of a μ heavy chain (μH or IgH μ) and a germ line-encoded surrogate light chain made of λ5 and VpreB (*9*). Intriguingly, the pre-BCR shapes the IgH repertoire and appears to select particular IgH chains, such as those that express an H-CDR3 with positively charged amino acid residues such as arginines (*10, 11*). Without any bias toward any particular signaling molecule, we carried out a series of experiments and discovered several lines of evidence to support that IgH is a critical DS receptor in a potential dual (DS•autoAg)-autoBCR signaling pathway and that DS is a potential master regulator of IgH.

## Results

### DS activity on NFS-25 cell proliferation and apoptosis

In our previous study, when primary mouse spleen cells were cultured with DS, CD5+ B-1 cells were found to be specifically stimulated by DS to proliferate (*1*). To test whether a similar effect holds for CD5+ pre-B cells, we cultured NFS-25 cells with DS, B-cell mitogen lipopolysaccharide (LPS), or medium only. Cell proliferation assays showed no significant difference between NFS-25 cells cultured with DS, LPS, or medium alone for 3 to 6 days. It is possible that these cells may have to be cultured with a much higher concentration of DS or LPS or for longer time periods to show detectable differences.

In a previous study, we had discovered that DS has strong affinity to apoptotic cells (*12*). We therefore examined whether this DS-affinity property is also true for NFS-25 cells. In typical NFS-25 cell cultures, about 5% to 8% of cells are observed to be apoptotic and stain positively with DS-Cy5 by flow cytometry. DS-Cy5 staining correlated well with annexin V staining but not with propidium iodide staining, consistent with our previous finding that DS has affinity to apoptotic cells. To further verify DS affinity in cell apoptosis, NFS-25 cell cultures were treated for different periods of time with various amounts of camptothecin (CPT), which triggers apoptosis by binding to the topoisomerase I and DNA complex in cells. As shown by flow cytometry, CPT induced apoptosis in a time- and dose-dependent manner (Supplemental Fig. 1). After overnight treatment, 10 μM CPT induced apoptosis in 26.8% of cells, while 50 μM CPT induced apoptosis in 41.3% cells, and all apoptotic cells were positively stained by DS-Cy5. These findings are consistent with our previously reported affinity of apoptotic cells to DS (*1*). We also tested whether DS could rescue CPR-induced apoptotic cells by co-culturing cells with DS and CPT, but DS did not increase the viability rates of CPT-treated cells.

### Possible DS affinity of CD5, CD72, CD19, PI3K, CD21, or Fas proteins

Since we hypothesize that DS•autoAg complexes engage dual signals, with autoAg binding to autoBCR and DS binding to another receptor, we began a search for unknown DS receptors by first screening prominent B-1 cell signaling markers. We extracted proteins from NFS-25 cells and fractionated them on DS affinity resins by step-wise elution with 0.2 M, 0.4 M, 0.6 M, and 1.0 M NaCl, corresponding to unbound or no, low, medium, and high DS affinities, respectively. Unfractionated and fractionated proteins were then separated in SDS-PAGE gels and blotted with antibodies to CD19, CD5, CD72, CD21, CD81, PI3K, Fas/CD95, and others (Fig. 1 and Supplemental Fig. 2).

**Fig. 1.**
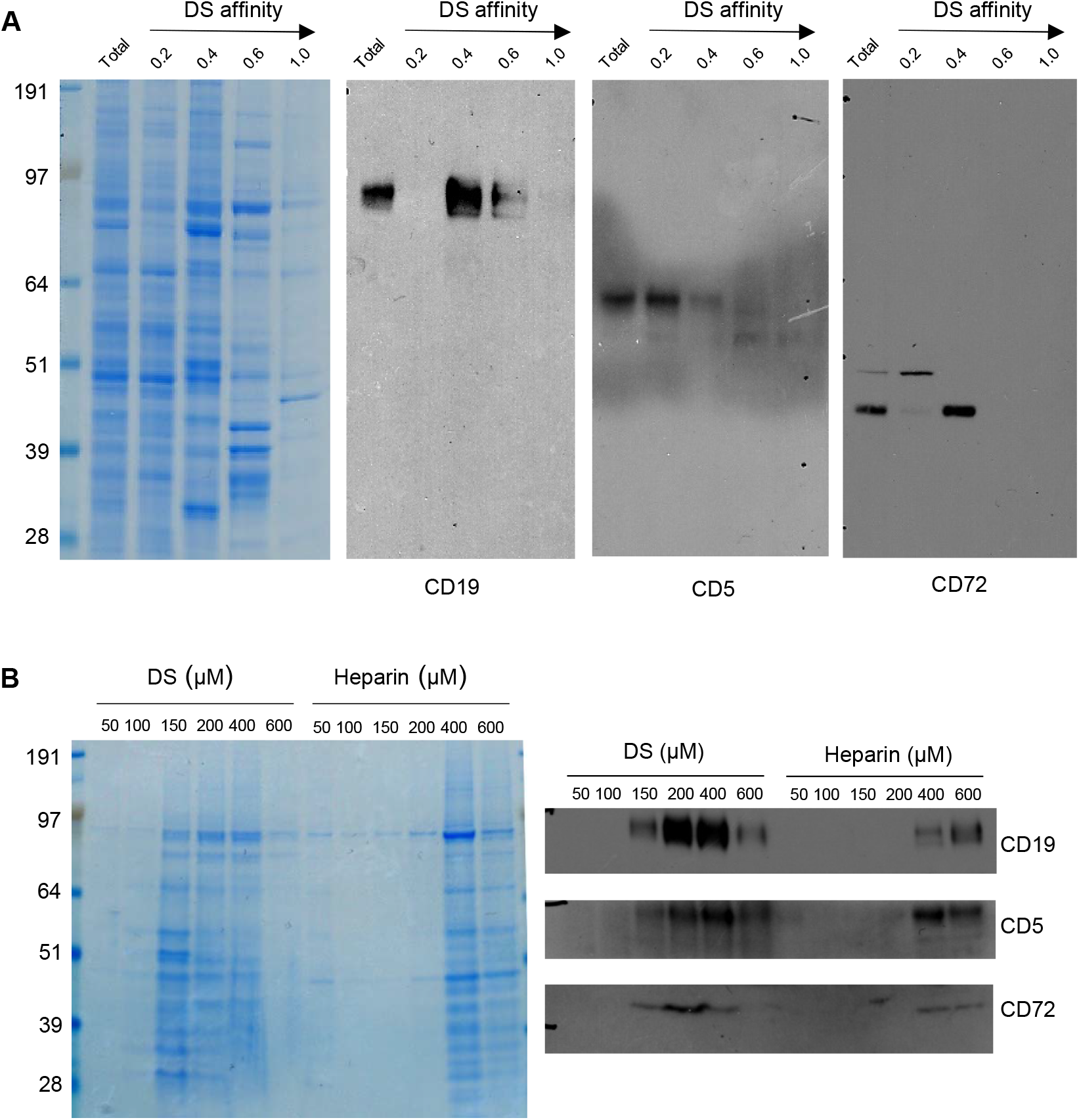
(**A**) Protein extracts from NFS-25 cells were fractionated with DS-affinity resins and blotted with specific antibodies. Protein lanes correspond to total unfractionated proteins (T) and fractions eluted with 0.2, 0.4, 0.6, and 1.0 M NaCl, respectively. (**B**) NFS-25 proteins loaded onto DS resins were eluted with step gradients of DS or heparin (left: Coomassie blue-stained SDS-PAGE) and blotted with specific antibodies (right), demonstrating competitive binding specificity between DS-resin-binding proteins and DS or heparin.

CD5 and CD19 are defining markers of B-1 cells, and CD72 is a known receptor for CD5. All three proteins were detected in the NFS-25 protein extracts as expected. After DS-affinity fractionation, CD19 was detected in the fractions eluted with 0.4 and 0.6 M NaCl, but not in the unbound (0.2 M fraction) or the 1.0 M fraction (Fig. 1A). CD5 molecules display a range of DS affinities, with the majority having no affinity but some having low affinity. However, two bands of smaller molecular size were also detected by anti-CD5 in fractions with medium and strong affinity. These findings suggest that CD5 may exist in multiple isoforms, and only some isoforms may associate with DS. CD72 was detected mostly in the 0.4 M fraction, although a higher molecular size version was present in the unbound fraction.

To ensure that the DS-affinity fractionation is indeed due to specific DS interaction but not non-specific binding to the resin, we carried out competitive elution with DS or heparin. NFS-25 protein extracts were loaded onto DS-Sepharose resins and eluted with DS or heparin (Fig. 1B). The majority of proteins bound to DS-Sepharose were eluted with DS at concentrations of 0.15, 0.2, or 0.4 M, whereas higher concentrations of heparin were needed. Western blotting verified that CD19, CD5, and CD72 were eluted with lower concentration of DS than heparin. These results suggest that proteins identified by our DS-affinity fractionation bind to DS more specifically than to heparin, a structurally similar polyanionic glycosaminoglycan.

We also examined a number of other markers as potential DS receptors (Supplemental Fig. 2). Phosphoinositide 3-kinase (PI3K) has been identified in the DS signaling pathway in mouse spleen cell cultures (*13, 14*). We therefore tested its affinity to DS. PI3K was detected in both the unbound and 0.4 M fractions. A smaller molecular size version was enriched in the unbound fraction, and a larger version was detected in the 0.4 M and 0.6 M fractions. Fas, also known as apoptosis antigen 1 or CD95, was identified in the unbound, 0.4 M, and 0.6 M fractions. Various bands with smaller and larger molecular sizes were also detected in the unbound, 0.4 M, and 0.6 M fractions. CD21 (or CR2) was detected in the unbound fraction.

Our previous study had shown that DS can also stimulate CD5+ cell proliferation in cells from CD19-deficient mice (*1*). The CD5 protein only partially co-localizes with DS (Supplemental Fig. 3). Although the above tested B cell signaling markers were detected in DS-binding factions and the results are consistent with our previous findings from mouse spleen B-1 cells (*1*), these proteins likely associate with DS indirectly via other molecules. Therefore, we continued our search for DS receptors by other means.

### DS trafficking in NFS-25 cells and accumulation in the ER

We tracked the fate of DS trafficking in cells by culturing NFS-25 cells with DS-AF568 and examined them by confocal fluorescence microscopy at various time points (Fig. 2). After 1 hour, DS appeared faintly on the surface of some cells. After 4 hours, DS association with small bodies, likely apoptotic cell bodies or dead cell debris, was observed. After 8 hours, some DS molecules were present within viable NFS-25 cells, starting to accumulate in an area adjacent to the nucleus. From 8 hours on, DS accumulation inside cells gradually increased. After 24 hours, DS accumulation in the same intracellular areas intensified and DS was also accumulating in the nucleus. After 48 hours, intracellular DS accumulation appeared to reach a maximum.

**Fig. 2.**
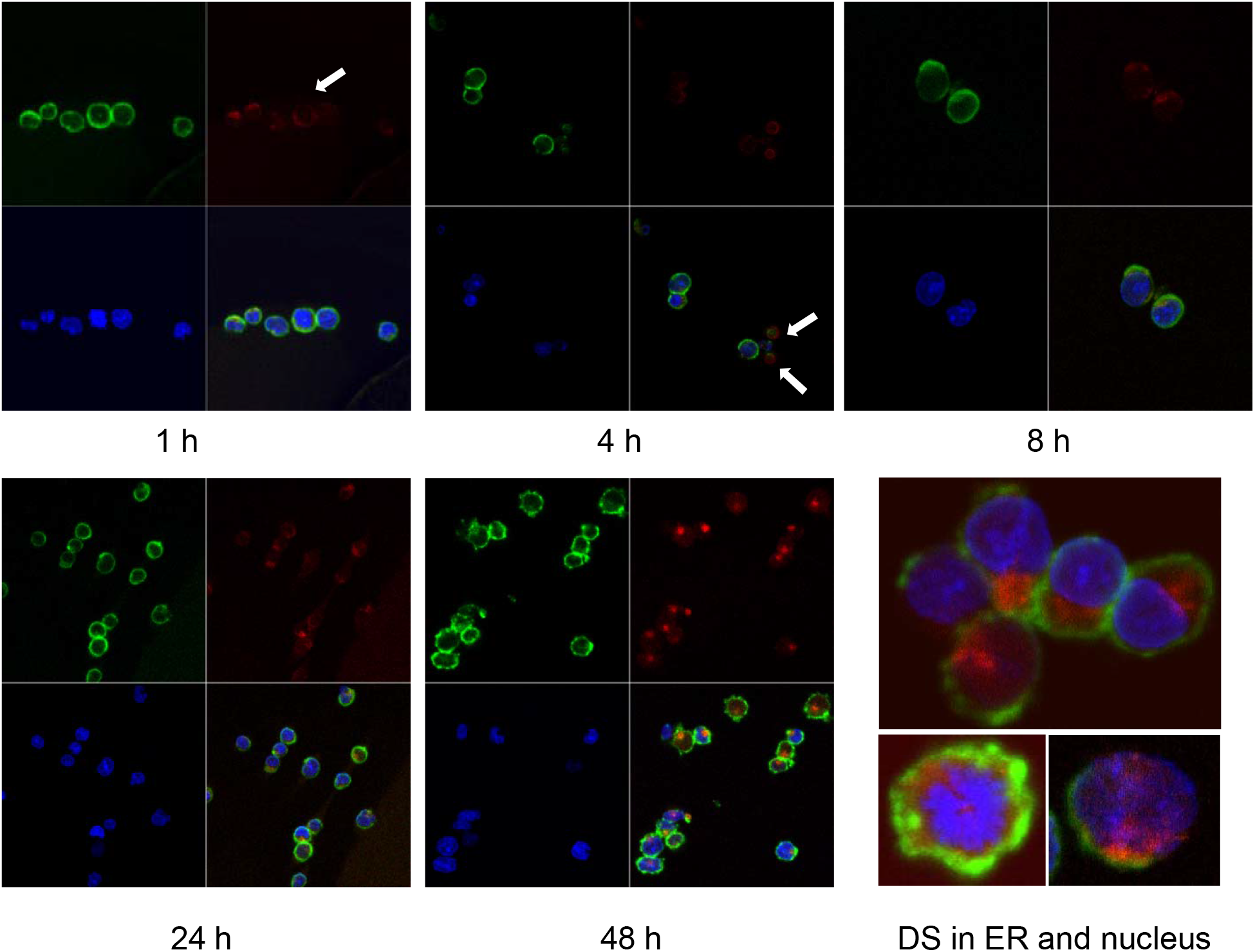
NFS-25 cells cultured with DS-AF568 (red) for various times were stained with anti-CD19 (green) and DAPI (blue). After 1 hour, trace amounts of DS was observed on some cell surfaces (white arrow). After 4 hours, DS association with small apoptotic cell bodies was observed (white arrows). From 8 hours on, DS starts to accumulate inside viable cells. At about 48 hours, intracellular DS reaches maximum intensity, with a majority in the ER and some also in the nucleus. Each panel shows four quadrants: green (upper left), red (upper right), blue (lower left), and merged (lower right) channels.

As shown in Fig. 2, DS did not randomly spread in the cytoplasm upon entering cells but rather accumulated for a significant amount of time in a specific compartment adjacent to the nucleus, consistent with the ER. Moreover, cells with accumulating DS appeared to be activated cells, which are slightly larger than the resting cells. Cells with nuclear dispersion of DS appeared even larger, suggesting nuclear uptake of DS in the synthesis or early mitotic phases of the cell cycle. To define the perinuclear DS-accumulating compartment, we stained the cells with antibodies for ER marker Grp78/BiP and Golgi marker Golgin97. Grp78 was found to co-localize with DS in the ER (Fig. 3A). To further verify Grp78 association with DS, we blotted NFS-25 proteins fractionated by DS affinity with anti-Grp78. Anti-Grp78 detected two strong bands and one faint band between 64 and 97 kDa molecular weight in the unfractionated NFS-25 protein extracts. The top band appeared predominantly in fractions with weak to strong DS affinity, whereas the bottom band appeared in all fractions. This result shows that Grp78 is abundantly expressed and that different forms can be associated with DS with weak to strong affinity (Fig. 3A).

**Fig. 3.**
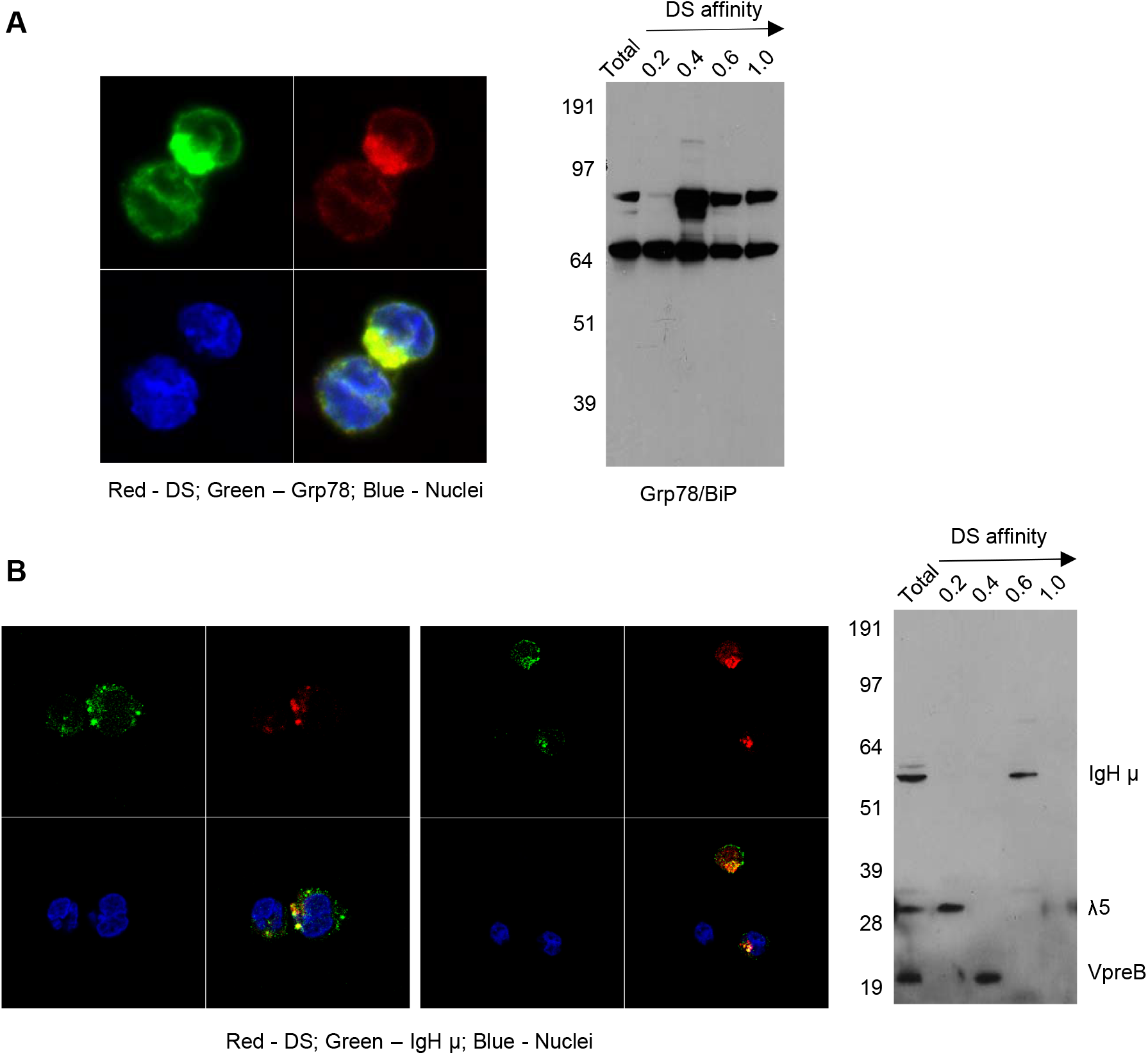
(**A**) Left: NFS-25 cells cultured with DS-AF568 and stained with anti-Grp78/BiP confirming the DS-accumulating compartment as the ER. Right: NFS-25 cell proteins fractionated by DS affinity and blotted with anti-Grp78/BiP. (**B**) Left: NFS-25 cells cultured with DS-AF568 and stained with anti-IgH μ showing co-localization on the cell surface and in the ER. Right: NFS-25 cell proteins fractionated by DS affinity and blotted with anti-preBCR, revealing the 3 components of preBCR, each of which has different DS affinity, with IgH μ having strongest affinity.

### Differential DS affinity of pre-BRC components IgH μ, λ5, and VpreB

The ER-resident protein Grp78 is also known as immunoglobulin heavy chain binding protein (BiP). We investigated whether immunoglobulins were involved in DS interactions in the ER. We stained NFS-25 cells cultured with DS-AF568 with anti-IgM heavy chain (IgH μ or μH), and, indeed, Grp78/BiP co-localized with DS in the ER (Fig. 3B). IgH μ also co-colocalized with DS in distinct clusters on the NFS-25 cell surface. In addition, IgH μ and DS co-localized faintly in the nucleus in a dispersed pattern.

We then examined whether DS stimulates Ig secretion. NFS-25 cells were cultured with either DS, anti-CD19, anti-CD5, anti-CD5+DS, anti-CD19+DS, anti-CD5+anti-CD19+DS, or medium only. IgH μ secretion was measured by ELISA. Anti-CD19 induced significant levels of IgH secretion. However, cells cultured with DS or anti-CD5 did not increase IgH secretion when compared to cells cultured with medium only. The addition of DS, anti-CD5, or anti-CD5+DS to anti-CD19 did also not enhance, but rather slightly reduced, IgH secretion when compared to cells cultured with anti-CD19 alone. It is possible that a DS-interacting site and CD5 may be in close proximity to CD19. Consequently, addition of DS or anti-CD5 would partially block and decrease anti-CD19-induced stimulation.

Pre-BCRs expressed in pre-B cells are formed by pairs of IgH μ chains and surrogate light chains composed of λ5 and VpreB proteins. In addition to IgH, we asked whether λ5 or VpreB also possess affinity to DS. We fractionated NFS-25 protein extracts and looked for pre-BCR components by blotting with a pre-BCR antibody that recognizes all three components (*15*). All three components were detected in the total unfractionated NFS-25 protein extract. Surprisingly, DS affinity neatly separated pre-BCR components into three distinct fractions: IgH chain appeared in the fraction eluting with 0.6 M NaCl, VpreB in the 0.4 M fraction, and λ5 in the 0.2 M fraction (Fig. 3B). This finding shows that the three components of the pre-BCR display differential affinity to DS.

These two complementary approaches, cell cultures with DS and co-localization microscopy and DS-affinity fractionation with Western blotting, demonstrate a strong interaction between DS and IgH μ protein. In summary, DS did not stimulate Ig secretion in NFS-25 cell but rather interacted with IgH μ on the cell surface, in the ER, and in the nucleus.

### Discovery of direct interaction between DS and GTF2I

Given that DS traffics from the cell surface to the ER and nucleus, there could be numerous additional molecules involved in the DS-pre-BCR signaling network. We, therefore, took an unbiased approach to identify DS receptors. NFS-25 cells were cultured with or without DS-biotin for 2 days, proteins were extracted from 50 million cells, and both insoluble and soluble portions were collected. Proteins were then pulled down with NeutrAvidin beads and compared side by side in SDS-PAGE gel lanes. No difference was detected in the insoluble portion, but two distinct bands at ~120 kDa molecular weight were detected in the soluble protein portion of cells cultured with DS-biotin (Fig. 4).

**Fig. 4.**
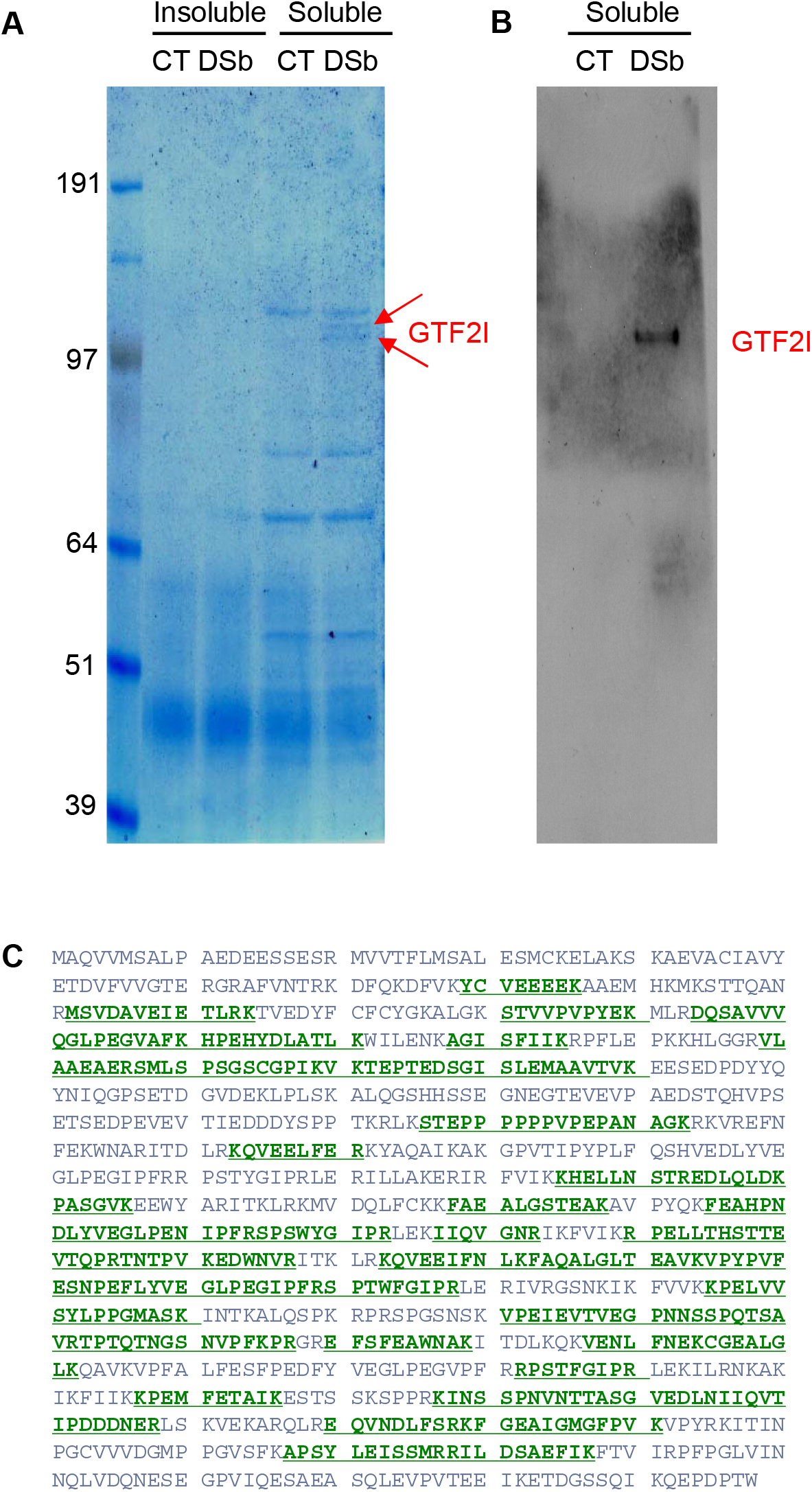
(**A**) Proteins extracted from NFS-25 cells cultured with DS-biotin (DSb) or without (CT, control) and analyzed by SDS-PAGE. The two arrows indicated two forms of GTF2I. (**B**) The identity of GTF2I is confirmed by blotting with anti-GTF2I. (**C**) Amino acid sequence of GTF2I (canonical isoform 1) with green regions highlighting peptides that were identified by mass spectrometric sequencing of the gel band indicated by the upper arrow in (**A**).

Both bands were sequenced by mass spectrometry. The top band identified a single protein with 46 peptide matches to GTF2I, which is impressive as mass spectrometry sequencing of protein bands from cellular extracts usually gives rise to many identifications. The bottom band also overwhelmingly identified GTF2I with 49 peptide matches (Fig. 4). A few additional proteins were also identified in the bottom band, including Uba1 (12 peptide matches), importin 7 (Ipo7, 4 peptide matches), ATP-citrate synthase (Acly, 3 peptide matches), kinesin heavy chain isoform 5A (Kif5a, 2 peptide matches), Ipo5 (2 peptide matches), and DNA ligase 1 (Lig1, 2 peptide matches). Western blotting also confirmed these protein bands as GTFT2I (general transcription factor II-I), a homolog of human protein TFII-I.

The GTF2I-encoding gene of humans or mouse contains six characteristic repeat motifs and at least 34 exons, and alternative splicing generates many transcript variants (*16*). Based on sequence BLAST, the peptides identified by mass spectrometry (covering 462 amino acid residues) did not align perfectly with any single known GTF2I isoform. But they did align best with mouse isoform X21 and aligned well with 30 other mouse isoforms, including 2, X6, X4, X14, 4, X5, X20, X12, 3, 1, X13, CRA-e, X3, X1, X2, X19, X11, X9, X18, CRA-g, 5, CRA-d, X8, X10, X16, X7, X15, or CRA-f.

### DS affinity protein network and IgH-associated ER complex

We reasoned that a potential DS receptor in a cellular molecular complex first needs to be accessible to allow DS association. Therefore, we attempted to tag accessible proteins in NFS-25 cells with amine-reactive biotin (sulfo-NHS-SS-biotin). Total proteins extracted from tagged cells were loaded onto DS-affinity resins and washed with 0.6 M NaCl, and those remaining tightly bound to DS were eluted with 1.0 M NaCl and collected. Biotin-tagged proteins were then pulled down with NeutrAvidin beads, separated in SDS-PAGE gels, and blotted with streptavidin-HRP for confirmation. Three unique bands were identified compared to those from mock experiments.

Mass spectrometry sequencing of these bands identified 40 proteins with a minimum of 6 peptide matches as a cut-off (Supplemental Table 1). These proteins are predominantly associated with the ER or the nucleus. Among these 40 proteins, 17 can be located to the ER, including BiP/GRp78 (Hspa5), Hsp90b1, Hsp90ab1, Ganab, VCP, Hspd1, Canx, Kpnb1, Hspa8, Pdia3, P4hb, Pdia4, Prkcsh, Rpn2, Hspa9, Cct8, and Cct3. 28 can be located to the nucleus, of which 8 both to the nucleus and the ER. The 20 exclusively nuclear proteins are Ncl, Hsp90aa1, Xrcc6, Xrcc5, Prmt5, Lmnb1, Eftud2, Supt16h, Supt5h, Mfap1b, Pkm, Hnrnpu, Ssrp1, Nasp, Mcm6, Actn4, Nap1l1, Eif3l, Pabpc1, and Tfrc. The remaining 3 proteins may be located to the cytoplasm or the cell membrane (Eif3b, Snx2, and Lck).

STRING protein-protein network analysis revealed that these 40 proteins have significantly more interactions than expected (Fig. 5). This network has 40 nodes and 182 edges (interactions) with average node degree of 9.1, average local clustering coefficient of 0.609, and PPI enrichment p-value of <1.0E-16, whereas a random set of 40 proteins is expected to have 42 edges. Based on KEGG pathway analysis, Protein Processing in the ER is the most significant for these proteins, with 13/161 counts in the network, 1.65 strength, and 1.80E-16 false discovery rate. Based on Gene Ontology analysis, Protein Folding is the most significant biological process from these proteins, with 12/153 counts in the network, 1.64 strength, and 1.53E-13 false discovery rate.

**Fig. 5.**
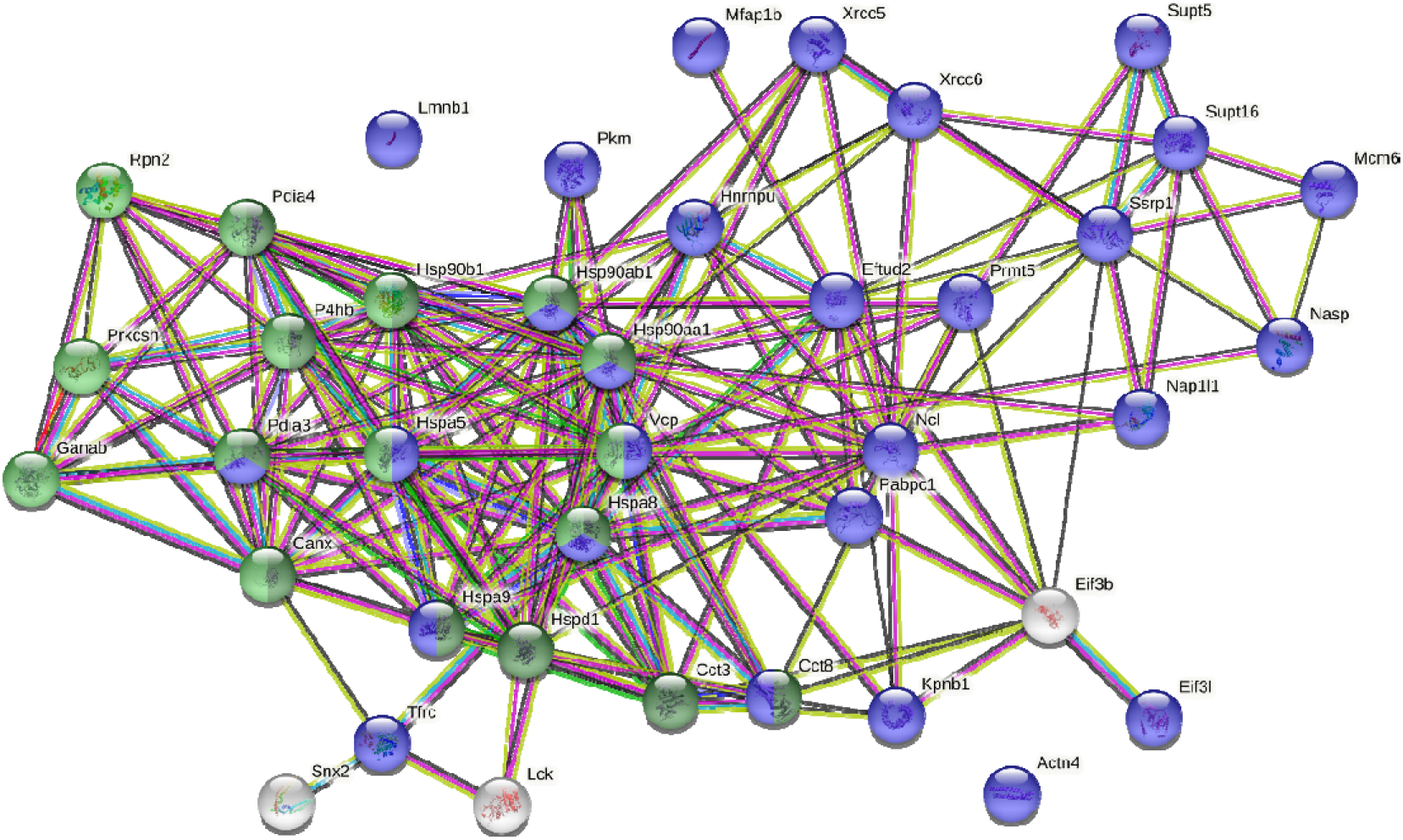
Protein-protein interaction network of the 40 biotin-tagged DS-binding proteins. Green: proteins involved in protein processing in the ER. Dark green: proteins involved in protein folding in the ER. Blue: nuclear proteins.

Pathway and process enrichment analysis by Metascape also revealed that these proteins are most significantly associated with Protein Folding (with Log_10_(P) value of −20.6) and Protein Processing in the ER (with Log_10_(P) value of −18.1). The Molecular Complex Detection (MCODE) algorithm applied to the densely connected network identified a single dominant complex among these DS-affine proteins, the IgH chain associated ER-localized multiprotein complex.

## Discussion

In this paper, we provide several lines of evidence to support DS as a master regulator of autoreactive B cell development at the pre-B stage via interaction with the IgH μ chain of the pre-BCR. Using CD5+ pre-B lymphoblast NSF-25 cells as a model, we show that DS interacts with IgH on the pre-B cell surface in distinct clusters, associates with an IgH-associated multiprotein complex in the ER, and recruits the general transcription factor GTF2I that may facilitate gene expression of *IgH* and other genes in the nucleus.

Based on these findings and our previous studies (*1-5, 17*), we propose an (DS•autoAg)-autoBCR dual-signal model for initiating autoreactive B-1 cell activation (Fig. 6): (i) DS and autoAgs from dying cells form non-covalent autoAg•DS complexes; (ii) upon encountering autoreactive B cells, an autoAg engages an autoBCR and binds to part of the variable domain (e.g., CDR1 and CDR2); (iii) because of its complexation with the autoAg, DS is brought into close contact with the autoBCR and binds to the IgH portion of the autoBCR (e.g., CDR3); (iv) (DS•autoAg)-autoBCR complexes internalize, and DS/IgH complexes accumulate in the ER where they may recruit IgH-associated protein processing complexes to facilitate folding and assembly of newly synthesized Ig; (v) somewhere along the way, DS binds GTF2I and transports it to nucleus, and in the nucleus, GTF2I upregulates transcription of the *IgH* locus and other gene targets. In summary, the intricate interactions of DS with Ig-associated complex in the ER and GTF2I may provide a positive feedback loop to upregulate IgH at gene and protein levels.

**Fig. 6.**
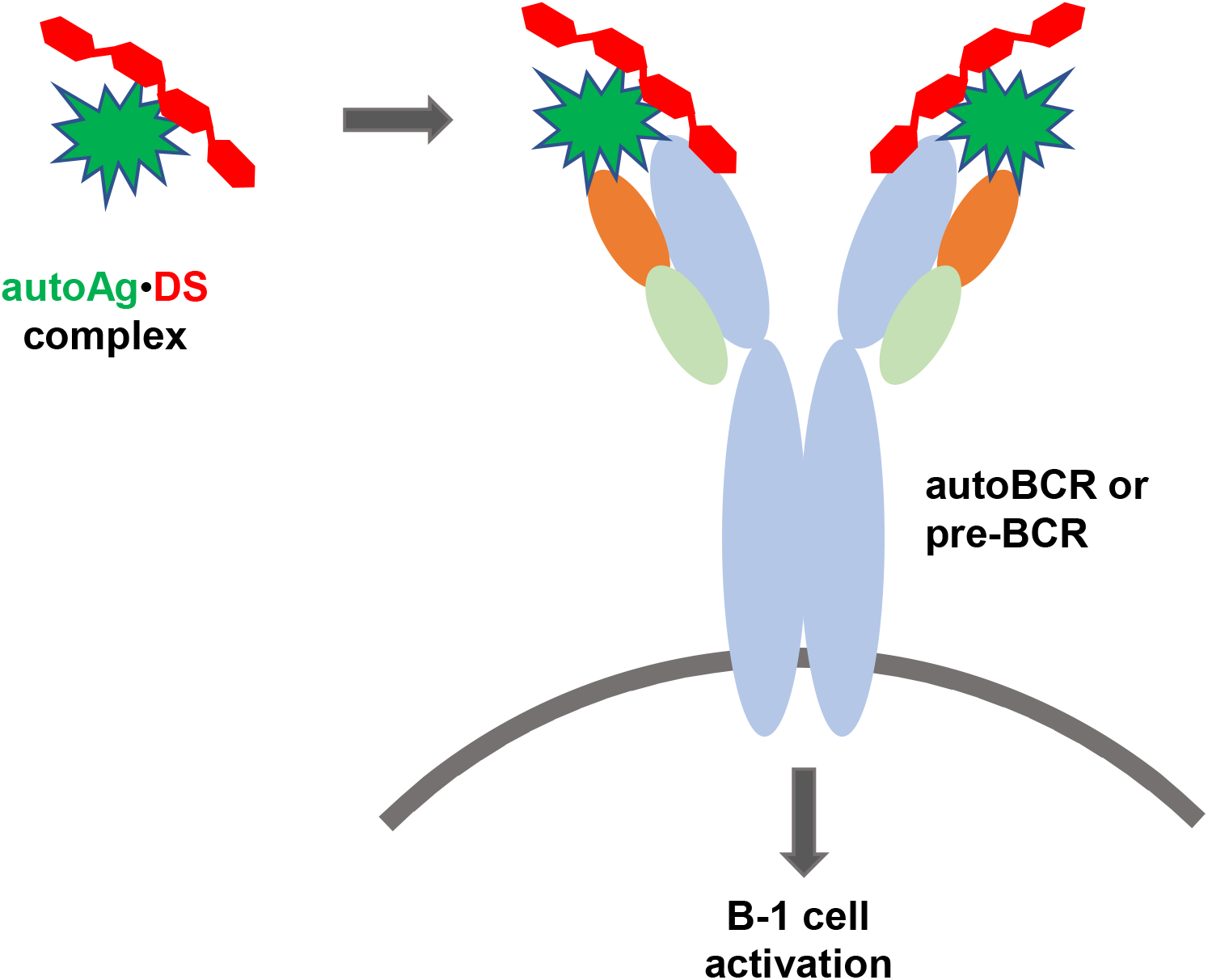
Proposed model of dual signaling by DS and autoAg in non-covalent (DS•autoAg)-autoBCR or (DS•autoAg)-pre-BCR complexes for activation of B-1 cells.

The IgH repertoire is generally thought to be positively selected at the pre-BCR checkpoint (*8*). The pre-BCR assesses the quality of IgH chains and tunes the repertoire by driving the preferential expansion and differentiation of cells with higher quality of IgH μ chains (*18*). In particular, the pre-BCR appears to favor IgH chains with a CDR3 region containing positively charged amino acid residues such as arginines (*10, 11*). Approximately 9% of in-frame IgH μ transcripts in human pro-B cells encode unusual IgH that can be expressed on the cell surface in the absence of surrogate or conventional light chains, and these IgH chains demonstrate preferential use of certain VH genes and an increased number of positively charged amino acid residues within the CDR3 region (*10*). Given that DS displays a high density repeating negative charges (on average, one carboxylate and one sulfate per repeating unit), we speculate that DS binds IgH at the CDR3 region.

Stromal cell-associated heparan sulfate, which is structurally similar to DS, has been proposed to be a potential pre-BCR ligand, as binding between pre-BCR and stromal cells could be specifically blocked by heparin, heparitinase, or a sulfate inhibitor (*19*). Moreover, the binding required the unique λ5 tail, which protrudes from the pre-BCR molecule at the same position where the CDR3 of a conventional light chain is located (*19*). We had tested heparan sulfate, heparin, and other glycosaminoglycans in our previous studies, but found DS to be the most potent glycosaminoglycan for stimulating B-1 cells (*1, 3*) and hence have focused on DS. Among the three components of the pre-BCR, our study found that IgH μ possesses strong DS affinity and λ5 weaker affinity (Fig. 3). It is possible that DS binds to a positively charged IgH CDR3 conformation that is enforced by the λ5 tail or conventional light chain CDR3.

Although synthesis and assembly rate of pre-BCR and BCR components are comparable, the pre-BCR is controlled by a highly efficient ER retention mechanism, which only allows exit of a few percent of the complexes from the ER (*20*). Furthermore, this retention mechanism is restricted to IgH and not selective for the surrogate light chain, thus appearing to be inherent to pre-B cells. Pre-B lymphocytes and hybridomas derived from them synthesize IgH in the absence of light chains, and BiP binds the CH1 domain of the free IgH and thus retains it in the

ER (*21, 22*). BiP also binds light chains shortly after their translation, and newly synthesized heavy and light chains interact sequentially with BiP and Grp94 in the ER (*23, 24*). BiP and other ER molecules strictly control the processing and assembly of immunoglobulins, and our current study identify DS as another critical player of the Ig-associated ER complex.

B cells synthesize large amounts of immunoglobulins, which makes these cells particularly sensitive to ER stress. When CHO cells were transfected with an IgH μ chain, expressed μ chain accumulated in the cells and formed stable complexes with BiP, and the synthesis of three ER stress proteins (BiP, Grp94, and ERp72) increased at both protein and mRNA levels (*25*). We have shown that DS accumulates in the ER and has affinity to these and other stress proteins.

Our study shows, for the first time, GTF2I protein as a direct DS-interacting partner in pre-B cells. The *GTF2I* gene is well known for its association with Williams-Beuren syndrome and supravalvular aortic stenosis. Amazingly, recent large-scale genome-wide association studies have identified close association of the *GTF2I* gene with various autoimmune diseases. Analysis of 556,134 autosomal SNPs in 542 cases and 1,050 controls followed by validation with 1,303 cases and 2,727 controls identified *GTF2I* at 7q11.23 (rs117026326) as a susceptibility locus for primary Sjögren syndrome (*26*). High-density genotyping of immune-related loci with replication in 4,478 SLE cases and 12,565 controls from six East Asian cohorts identified *GTF2IRD1-GTF2I* at 7q11.23 to be the most significant locus associated with SLE (*27*). Variants at the *GTF2IRD1-GTF2I* locus have been identified as a susceptibility locus shared by SLE (*27*), primary Sjögren syndrome (*26*), rheumatoid arthritis (*28*), and myasthenia gravis (*29*). Single nucleotide polymorphisms (SNPs) located in the *GTF2I-NCF1* region (rs73366469 (*GTF2I*), rs117026326 (*GTF2I*), rs80346167(*GTF2IRD1*) and rs201802880 (*NCF1*)) have strong association with SLE, Sjögren syndrome, rheumatoid arthritis, and ANCA-associated vasculitis (*30*). SNPs at rs117026326 are also associated with neuromyelitis optica spectrum disorder (*31*). *GTF2I* gene mutations occur at significantly higher frequencies in thymoma patients with myasthenia gravis than those without (*29*).

GTF2I proteins are unusual transcriptional regulators that function as both basal and signal-induced transcription factors. Among its related pathways are the BCR signaling pathway and RNA polymerase II transcription initiation and promoter clearance. GTF2I protein interacts with Btk (Bruton’s tyrosine kinase), ARID3A (AT-rich interaction domain), HDAC3 (histone deacetylase 3), and others (*16*). GTF2I controls B cell proliferation by regulating both nuclear translocation of c-Rel and DNA-binding activity of NF-kappaB (*32*).

GTF2I interacts directly with B cell-specific coactivator OCA-B and regulates IgH gene transcription by facilitating enhancer-promoter communication (*33*). GTF2I is also required for induction of IgH gene transcription through transcription factor Bright (B cell regulator of immunoglobulin heavy chain transcription) (*34*). GTF2I, Btk, and Bright function as a three-component protein complex at the *IgH* gene locus. GTF2I and Btk exist in a complex before BCR engagement, and in response to BCR crosslinking, GTF2I becomes transiently phosphorylated on Tyr248, Tyr357, and Tyr462 (*35, 36*). Multiple kinases can independently target GTF2I via distinct signaling pathways. In addition to Btk (*37*), GTF2I undergoes c-Src-dependent phosphorylation on Tyr248 and Tyr611 and translocates from the cytoplasm to the nucleus (*38*).

GTF2I is required for optimal induction of GRP78/BiP and others in the ER stress response (*39, 40*). When cells experience ER stress, the transcription of a family of glucose-regulated protein (*GRP*) genes which encode various ER chaperones is induced. The *GRP* promoters contain multiple copies of ER stress response element (ERSE), consisting of a unique CCAAT(N_9_)CCACG tripartite structure with N_9_ being a GC-rich sequence of 9 bp that is conserved across species (*40*). GTF2I isoforms bind directly to the ERSEs of grp78 and ERp72 (PDIA4) promoters, and the stimulation of ERSE-mediated transcription by GTF2I requires consensus tyrosine phosphorylation sites on GTF2I and the ERSE GGC sequence motif (*40*). GTF2I binds the Grp78 gene promoter and in turn regulates Grp78 transcription and synthesis (*41*). In our study, we identified GRP78 (BiP), GRP94 (Hsp90b1) and ERp72 (PDIA4), as well as a number of other ER stress chaperones such as GRP75 (Hspa9), Hsp90ab1, Hspd1, Hspa8, and Hsp90aa1 (Fig. 5 and Supplemental Table 1). It is possible that GTF2I regulates the transcription of these proteins.

Our study also identified 20 nuclear proteins, which are likely involved in GTF2I associated gene transcription activities. In addition, LCK is identified as a DS-associated protein. Although typically regarded as a T-cell specific tyrosine kinase, LCK is an important mediator of BCR signaling in B-cell chronic lymphocyte leukemia (B-CLL) and found at significant levels in CD5+ B-1 cells (*12, 42*). In mice, LCK is also expressed by CD5+ B-1 cells and involved in modulating BCR signal transduction in B-1 cells (*43, 44*).

In our previous studies, we have demonstrated that DS has strong affinity to over 200 autoAgs and proposed that autoAgs and DS form macromolecular complexes and cooperate to stimulate autoreactive B cells (*1, 2, 4, 5*). Therefore, we have been searching for DS receptors in autoreactive B cells. In this study, we identified IgH μ of the pre-BCR as a novel DS receptor. Pre-BCRs resemble polyreactive autoBCRs in their capability of recognizing multiple autoAgs (*7*). Autoreactive B-1 cells differ from conventional B-2 cells in their responses to BCR signaling. While binding of foreign antigens to BCRs on B-2 cells leads to cell activation, repeated encounters with autoAgs switch B-1 cells either into an anergic state or lead to apoptosis.

We propose a model of (DS•autoAg)-preBCR or (DS•autoAg)-autoBCR dual signaling that activates autoreactive B cells (Fig. 6). In this model, both autoAg and DS cooperate and bind simultaneously to different parts of the autoBCR. Once internalized with the BCR complex, DS recruits a network of additional partners that aids initial B-1 cell activation to proceed to the next stages of B cell development.

In summary, this study provides several lines of evidence that support DS as a potential master regulator of IgH in pre-B cells. DS interacts directly with IgH on the cell surface, with IgH-associated protein processing complexes in the ER, and with the GTF2I transcription factor that is required for *IgH* transcription. Through its affinity with autoAgs and its control of IgH, DS emerges as a potential key player in the development of autoreactive B cells and autoimmunity.

## Materials and Methods

### Cell culture

NFS-25 C-3 cells were purchased from the ATCC (CRL-1695) and cultured in complete DMEM supplemented with 10% fetal bovine serum and a penicillin-streptomycin-glutamine mixture (Invitrogen). For various experiments, cells were cultured in the medium with the addition of DS or heparin (10, 50,100, or 500 μg/ml), 10 μg/ml of LPS, anti-CD5 (2, 5, 10 μg/ml), anti-IgM (2, 5, 10 μg/ml), or combinations for 3-6 days. Cell proliferation was measured by MTT assays. For apoptosis induction, cells were cultured with 5, 10, 20, 50 μM camptothecin (CPT) for various times. Cell apoptosis was monitored by Annexin V-FITC and PI staining and flow cytometry.

### Protein extraction

Approximately 200 million NFS-25 cells in 200 mL cell culture medium were harvested and washed twice with PBS. They were suspended in 50 mM phosphate buffer (pH 7.4) with Roche Complete Mini protease inhibitor cocktail. Cells were homogenized on ice, followed by sonication on ice for 5 min. The homogenate was centrifuged, and the supernatant was considered the total protein extract and collected. Protein concentrations were measured with the RC DC Protein Assay (Bio-Rad).

### DS-affinity fractionation

DS-Sepharose resins were prepared by covalently linking DS (Sigma-Aldrich) to EAH Sepharose 4B resins (GE Healthcare) as previously described (*1, 2*). Aliquots of about 5.5 mg of protein extract in 1 ml of 10 mM phosphate buffer (buffer A, pH 7.4) was mixed with 0.35 ml of DS-Sepharose resins and incubated at 4 °C for 3 hours. After removing the buffer, the resins were washed three times with 1 ml buffer A to remove unbound proteins. Proteins bound to resins were then eluted stepwise and sequentially with 0.2, 0.4, 0.6, 1.0 M NaCl in buffer A, corresponding to proteins with no, low, medium, to high DS affinity, respectively. For each salt elution, resins were mixed with 1.0 ml of elution buffer for 5 min, then the eluate was collected, and each elution step was repeated three times. Eluted fractions were desalted and concentrated in 5 kDa cut-off Vivaspin centrifugal filters (Sartorius).

### Competitive elution with DS and heparin

Proteins were extracted from NFS-25 cells and loaded onto DS-Sepharose resins as described above. After washing with 10 mM phosphate buffer three times, the resins were divided into two aliquots. In one aliquot, proteins were eluted sequentially with 0.2 ml of 0.05, 0.1, 0.15, 0.2, 0.4, and 0.6 mM DS. In the other aliquot, proteins were eluted sequentially with 0.2 ml of 0.05, 0.1, 0.15, 0.2, 0.4, and 0.6 mM heparin. Samples of eluted fractions were analyzed by SDS-PAGE, stained with Coomassie blue, and blotted with anti-CD19, anti-CD5, and anti-CD72.

### SDS-PAGE and Western blotting

SDS-PAGE analysis of proteins was performed with 4-12% NuPAGE Bis-Tris gels (Invitrogen) with MOPS running buffer and staining with Bio-Safe Coomassie G250 (Bio-Rad). Aliquots of ~5 μg of proteins were loaded per gel lane. Proteins in gels were transferred onto PVDF membranes, blocked with TBS (pH 7.4) containing 2% BSA, 3% casein or skim milk, and 0.5% Tween 20 at 4 °C overnight. They were incubated with primary antibodies at 25 °C for 1 h. Primary antibodies tested include anti-CD19 (eBioscience), anti-CD5 (sc-6985, Santa Cruz Biotech), anti-CD72 (sc-25265), anti-CD21 (sc-7027), anti-CD81 (sc-9158), anti-PI3K (sc-31969), anti-Grp78 (Abcam), anti-Golgin97 (Abcam), biotin anti-CD95 (Fas/APO-1, eBioscience), or biotin rat anti-mouse preBCR (BD Biosciences). Membranes were washed thrice with TBS containing 0.5% Tween 20, then incubated with appropriate secondary antibodies conjugated with horseradish peroxidase in blocking buffer at 25 °C for 1 h, and developed with ECL substrate. Secondary antibodies used include goat anti-rabbit IgG-HRP, anti-goat IgG-HRP, or streptavidin-HRP (BioLegend).

### DS-biotin isolation of GTF2I

DS-biotin was synthesized as previously described (*1*). NFS-25 cells were cultured with and without 20 μg/ml DS-biotin in complete DMEM medium for 3 days. Proteins were extracted from 50 million cells using the MEM-PER extraction kit (Thermo Fisher). Soluble proteins (1.5 ml) and insoluble proteins were separated, and to each tube was added 1.5 ml of 10 mM phosphate buffer and 0.06 ml of NeutrAvidin agarose beads (Pierce). The mixtures were incubated at 4 °C for 1 h. After washing the beads with PBS four times, bound proteins were released by boiling with SDS-PAGE sample buffer at 100 °C for 10 min. Proteins from cells cultured with or without DS-biotin were compared side by side by SDS-PAGE analysis. Bands only present in samples from cells cultured with DS-biotin were selected and sequenced by mass spectrometry. In duplicate gels, proteins were transferred onto PVDF membrane, blotted with anti-GTF2I (sc-9943, Santa Cruz Biotech) at 1:1000 dilution, and detected with rabbit anti-goat IgG-HRP at 1:2000 dilution and ECL substrate.

### Cellular protein biotinylation and DS-affinity isolation

NFS-25 cells cultured in complete DMEM were harvested, washed with ice-cold PBS three times, and resuspended at a concentration of 25,000 cells/ml in PBS. To each 1 ml of cells was added 80 μl of 10 mM sulfo-NHS-SS-biotin (Pierce) in H_2_O and incubated at room temperature for 30 min. After the reaction, the cells were washed three times with ice-cold PBS and lysed by homogenization. Cell lysates were incubated with 1 ml of DS-Sepharose resins, and unbound to weakly-bound proteins were washed off extensively with 0.4 M NaCl in 10 mM phosphate buffer. Tightly bound proteins were eluted with 1.0 M NaCl in 10 mM phosphate buffer. The eluate was incubated with 0.2 ml of streptavidin agarose resins (Pierce) at 4 °C for 1 h. The resins were washed four times with PBS, and bound proteins were eluted by boiling with 0.2 mL SDS-PAGE buffer for 10 min. The eluted proteins and controls were analyzed by SDS-PAGE, transferred to PVDF membranes, and blotted with streptavidin-HRP to reveal biotinylated proteins. Biotinylated protein gel bands were selected and sequenced by the Taplin mass spectrometry facility at Harvard Medical School (Boston).

### Cell proliferation and Ig μ measurement by ELISA

IgM secreted into the cell culture supernatant was measured by ELISA. Duplicate wells in 96-well plates were coated with rabbit anti-mouse F(ab’)_2_ in 0.1 M sodium carbonate buffer (pH 9.4) at 4 °C overnight. After washing with 0.05% Tween 20 in PBS three times and blocking with 1% BSA at 37 °C for 1 hour, 100 μl of cell culture supernatant was added to each well and incubated at 37 °C for 1 hour. IgH μ heavy chains were detected with goat anti-mouse μ conjugated to alkaline phosphatase (SouthernBiotech) and p-nitrophenyl phosphate substrate.

### Flow cytometry

Cultured cells were collected by centrifugation and washed twice with a staining buffer that contained 1% bovine serum albumin in PBS (pH 7.2). About 4 million cells were suspended in 0.8 ml of the staining buffer and split into eight 0.1 ml aliquots, ~0.5 million cells each. Aliquots were stained with 1-3 μg of anti-CD19-FITC, PE anti-CD5-PE, DS-Cy5, or a combination of the three by incubating in the dark at 25 °C for 1 hour. In another experiment, cells were stained with 1 μg of anti-IgM-FITC, anti-IgD-FITC, DS-Cy5, or combinations. Stained cells were washed three times with 0.5 ml of the staining buffer, and analyzed with a BD FACS Calibur flow cytometer.

### Fluorescence confocal microscopy

NFS-25 cells were cultured with 10 μg/ml of DS-AF568 for 1-6 days. Cells were fixed with 1% formalin in PBS at 25 °C for 15 min and washed twice with PBS. Cells were stained in PBS containing 1% BSA or 10% FBS with various antibodies at 4 °C for 18 h. Antibodies used include: anti-CD19-FTIC, anti-CD5-FITC, anti-Grp78-AF488, and anti-μ-FITC (SouthernBiotech). Cell nuclei were stained with 50 nM DAPI at 25 °C for 15 min. Images were taken with a Zeiss LSM 510 confocal microscope.

## Acknowledgements and Funding Statement

We thank Ross Tomaino and the Taplin Biological Mass Spectrometry facility of Harvard Medical School for expert service with protein sequencing. This work was partially supported by Curandis and the NIH of the United States. MHR acknowledges NCI R21 CA251992, funding from a Cycle for Survival Equinox innovation grant, and the MSKCC NIH/NCI Cancer Center Support Grant P30 CA008748.

## Author Contributions

JML carried out the experiments. JHR assisted with experiments. MHR consulted on the study and edited the manuscript. JYW directed the study, analyzed the data, and wrote the manuscript. All authors have proofread the paper.

## Competing Interest Statement

JML and JHR declare no competing interest. MHR is on the scientific advisory boards of Trans-Hit, Proscia, and Universal DX. JYW is the founder of Curandis. MP Biochemicals is the current employer of JHR. None of these companies had influence on the design, interpretation, or decision to publish of this study.

## Classification

Autoimmunity

## Field codes

Precursor BCR, Dermatan Sulfate, Ig Heavy Chain, GTF2I, BiP

---

**Supplemental Table 1.**
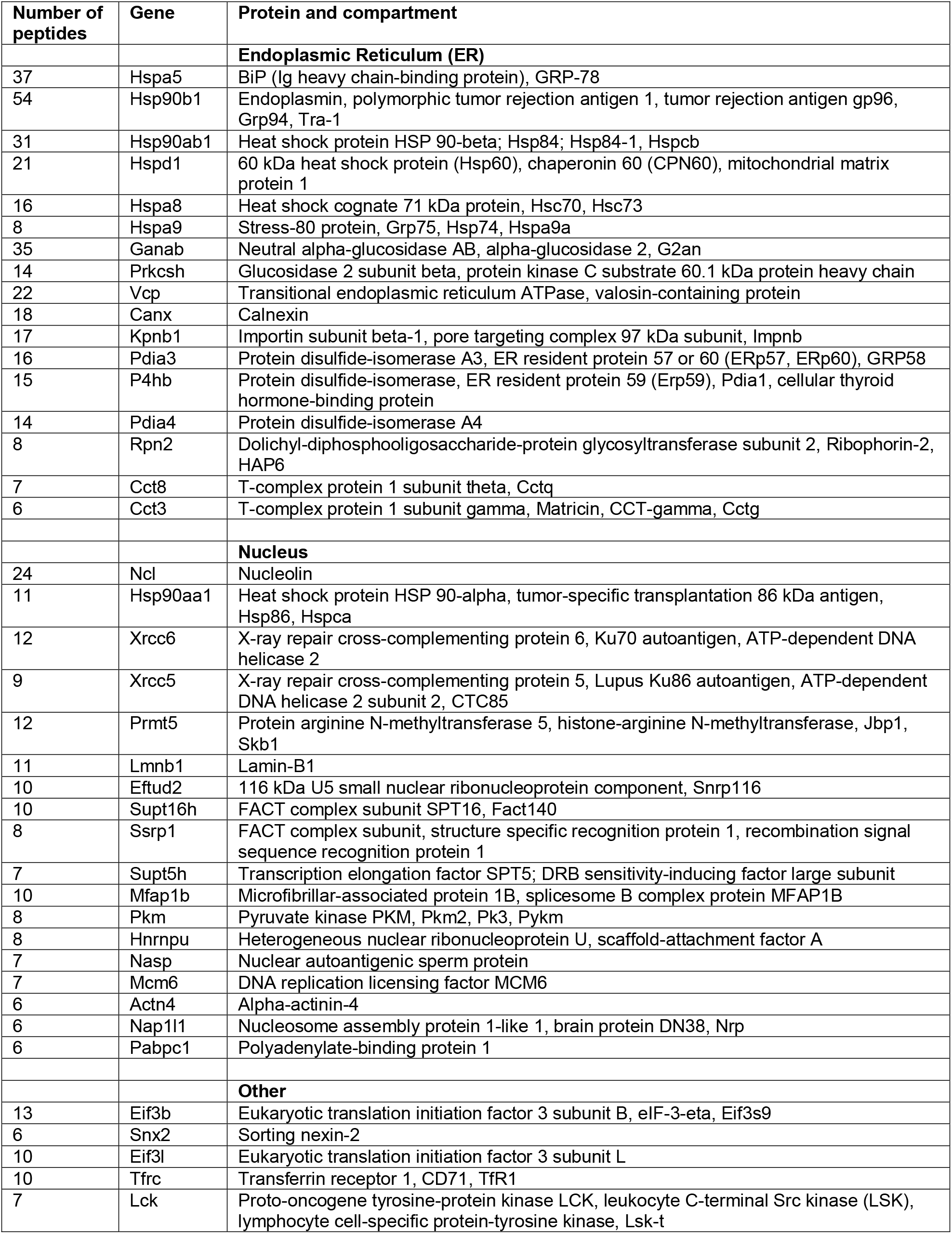
Proteins identified from NSF-25 cells with strong DS affinity.

**Supplemental Fig. 1.**
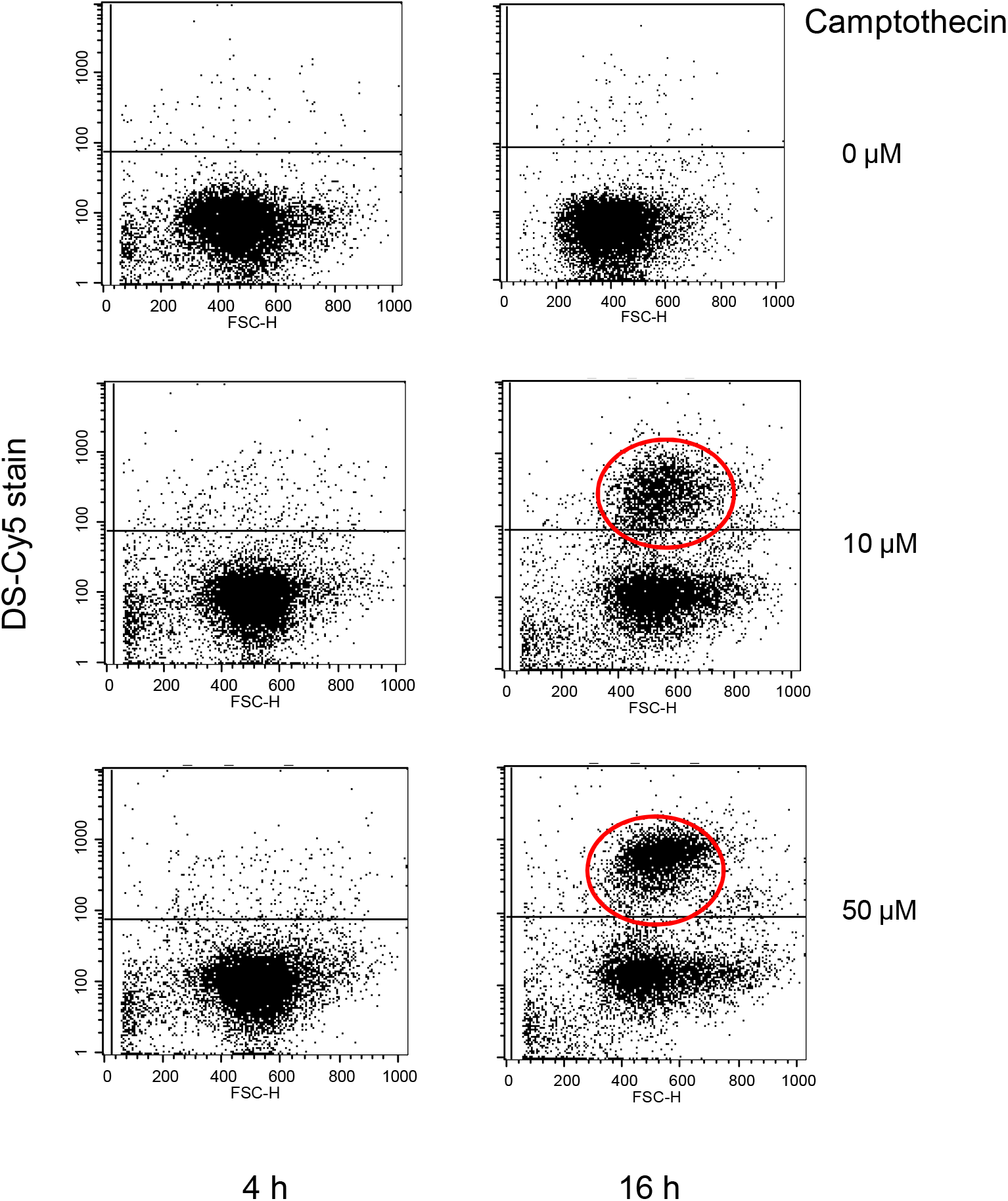
NFS-25 cells cultured with camptothecin and stained with DS-Cy5 to demonstrate the binding of DS to apoptotic cells (red circles).

**Supplemental Fig. 2.**
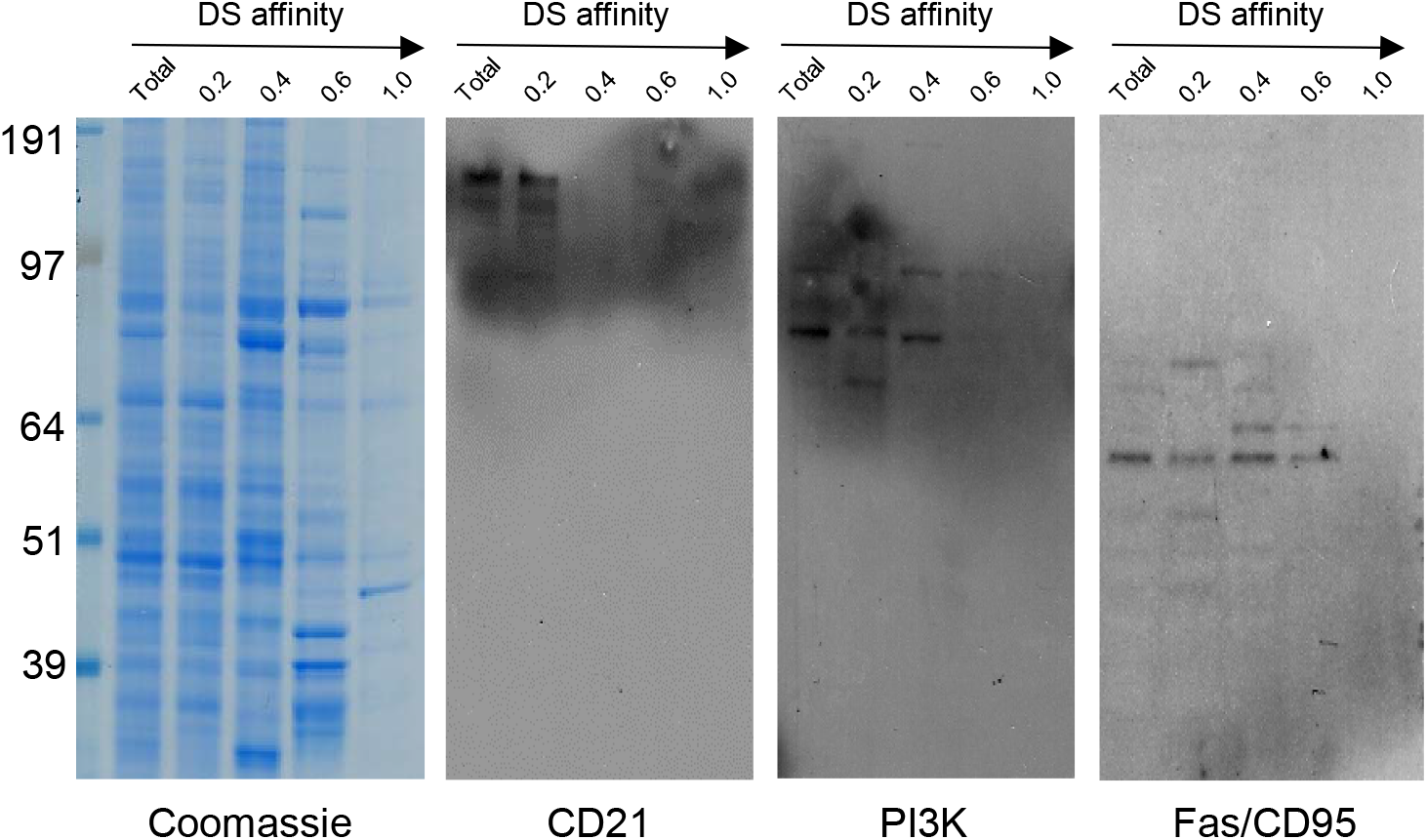
NFS-25 cell proteins were fractionated by increasing DS affinity (left to right) and blotted with anti-CD21, anti-PI3K, and anti-Fas (CD95).

**Supplemental Fig. 3.**
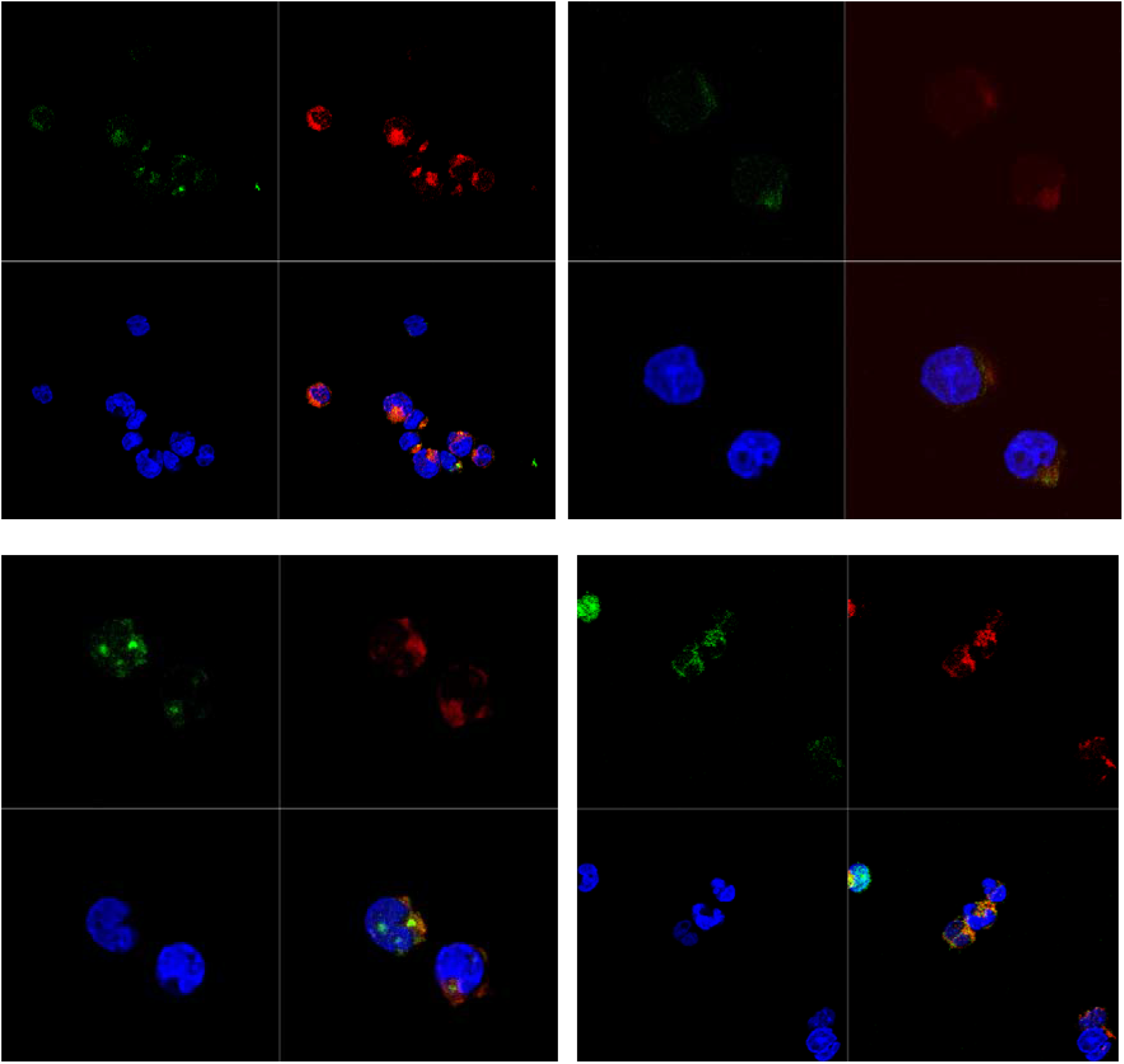
NFS-25 cells cultured with DS-AF568 (red) and stained with anti-CD5 (green) and DAPI. DS and CD5 only partially co-localize. Each panel shows four quadrants: green (upper left), red (upper right), blue (lower left), and merged (lower right) channels.

